# Impact of selective digestive decontamination on the pangenome composition of ESBL-*E. coli*

**DOI:** 10.1101/2025.01.14.632990

**Authors:** Julian A. Paganini, A. C. Schürch, Jelle Scharringa, M. J. M. Bonten, R. J. L. Willems, N. L. Plantinga

## Abstract

In Dutch ICUs patients receive selective digestive decontamination (SDD) as a prophylactic antimicrobial treatment to prevent colonisation with potentially pathogenic microorganisms and subsequent infections, with a beneficial effect on 28-day mortality. In the R-GNOSIS ICU study, conducted outside of The Netherlands, SDD consisted of a mix of an oropharyngeal paste and a gastric suspension containing colistin, tobramycin and nystatin. These topical antimicrobial agents aim to target aerobic Gram-negative bacteria, *S. aureus* and yeast. SDD improves patient outcome, but its effects on the resistome and pangenome of potentially pathongenic microorganisms have not been extensively studied. In this work, we compared 129 genomes of *E. coli* isolates from patients that received SDD and patients that did not receive SDD, but standard care only (baseline patients) in five ICUs located across three European countries (R-GNOSIS ICU study). We found that the overall pangenome compositions of *E. coli* recovered from both patient groups were highly similar. Variations in the accessory genome were strongly associated with the phylogeny of isolates but not with the use of SDD. Similarly, the plasmidome variations were not explained by treatment, but rather by the interaction between ICU location and phylogroup. One antibiotic resistance gene, which confers resistance to tobramycin, was found to be more prevalent in SDD patients. This gene was flanked by IS26 elements and significantly co-occurred with blaCTX-M-15 in various plasmid backbones and chromosomal contexts. Notably, no mcr genes coding for colistin resistance were detected.

## Introduction

In the Netherlands, patients admitted to the ICU and undergoing mechanical ventilation receive selective digestive decontamination (SDD) as a prophylactic treatment to prevent colonisation with potentially pathogenic microorganisms (PPMOs). SDD consist of a mix of topical antibiotics (tobramycin, colistin and amphotericin B) targeting aerobic gram-negative bacteria (GNB), *Staphylococcus aureus* and yeast, but leaving the anaerobic flora intact. SDD is administered as an oropharyngeal paste and as a solution through the nasogastric tube. Additionally, a 4-day course of an intravenous cephalosporin (cefotaxime or ceftriaxone) is also provided, to treat any incubating infection at the time of ICU admission ^1–3^. A variation of SDD, named Selective Oropharyngeal Decontamination (SOD), consisting only of the oropharyngeal paste, is administered in some ICUs as an alternative to SDD ^4^. In the Netherlands, where the prevalence of antibiotic resistance is low ^5^, SDD was associated with improved patient outcome in comparison to standard care, with reduced mortality, lengths of ICU stay and a lower incidence of ICU-acquired bacteremia ^6–9^.

The R-GNOSIS ICU study, conducted between 2013 and 2017, compared the effectiveness of SDD, SOD, chlorhexidine 1% mouthwash and standard care (baseline) in thirteen ICUs located in six European countries with medium to high prevalence of antibiotic resistance (defined as having an extended-spectrum β-lactamase -ESBL-prevalence of at least 5% of amongst ICU-acquired bacteremia with Enterobacteriaceae) ^10^. In this cluster-randomised trial, each treatment was applied during six months in the entire ward (randomised order), to patients with expected length of mechanical ventilation of at least 24h. An important modification of the SDD regime in this study was the absence of the 4-day course of intravenous cephalosporin.

One concern regarding the application of SDD is the exertion of antibiotic pressure in ICUs, which already have the highest levels of antimicrobial use within hospitals ^11^. In contrast to this, multiple studies suggest that the use of SDD is associated with a decrease in incidence of colonization and infection with antimicrobial resistant microorganisms, both in settings with high and low prevalence of resistance ^12–14^. On the other hand, a study based on metagenomic data seems to indicate that resistance genes to three classes of antibiotics, namely aminoglycosides, macrolides and tetracyclines, are more abundant in the gastrointestinal tract of SDD treated patients when compared to healthy individuals ^15^. Moreover, a separate study concluded that during the application of SDD in a single ICU patient, the burden of two aminoglycoside resistance genes seemed to increase in culturable anaerobic commensal bacteria ^16^. Furthermore, these two genes appear to be located on mobile genetic elements (MGEs), increasing the risk of horizontal gene transfer (HGT) from anaerobic bacteria to PPMOs. In addition to the potential selective effects on resistance genes, studies also indicate that SDD treatment alters the gut microbiome composition of ICU patients ^17,18^. These ecological changes of the microbiota may also affect the composition of PPMOs, such as *Escherichia coli*, populating the intestinal tract of patients receiving SDD. Therefore, we hypothesise that SDD treatment shapes the pangenome composition of *E. coli*, including the development of resistance.

To address this question, we have sequenced the genomes of 129 *E. coli* isolates from the R-GNOSIS ICU study. These isolates were obtained from patients that received either SDD (n=63) or did not receive SDD (n=69, baseline) in five different ICUs located in Spain, Belgium and the UK. We explored the population structure of these isolates and compared their accessory genome content, plasmidomes and resistomes to determine if SDD leads to the selection of specific genomic features of *E. coli*.

## Materials and Methods

### R-GNOSIS ICU study and selection of isolates for whole genome sequencing

The R-GNOSIS ICU study was conducted between December 2013 and May 2017 in 13 ICUs from six European countries. A detailed description of the study’s aims and methods can be found in ^10^. Briefly, to monitor the effect of SDD on colonization with Gram-negative bacteria, surveillance samples were taken twice weekly from the rectum and respiratory tract of all patients included in the study (n=8,496). Samples were inoculated on ESBL selective media (Biomerieux®) and in case of growth, phenotypic susceptibility testing was performed according to local standard operating procedures (for colistin susceptibility testing, E-tests were provided) ^10^. Clinical blood and respiratory samples were obtained at discretion of the clinician and processed according to local laboratory protocols. From these cultures, unique highly-resistant microorganisms were stored for whole genome sequencing (WGS) according to the following rule: one isolate per patient, per body site, per species, with a unique phenotypic resistance pattern.

We submitted for WGS all stored *E. coli* isolates from the SDD and baseline periods of the five hospitals with most stored isolates (AN, PS, UZ, LB and CD). Baseline isolates were defined as those collected from patients who did not receive SDD treatment during the study. Isolates were only included if they were collected from day 2 of inclusion onwards (with the date of study enrollment being day 0), to ensure sufficient exposure to the antimicrobials used in SDD. Metadata associated with sequenced isolates can be found in Supplementary Data 1.

### Whole Genome Sequencing

Selected isolates were sequenced using Illumina MiSeq, with a Nextera XT pair-end kit (2 x 150bp). Short reads were quality trimmed with trim-galore (v0.6.6) (https://github.com/FelixKrueger/TrimGalore). Assembly of genomes was performed with Unicycler (v0.4.9) ^19^.

### Pangenome analysis

Genomes were first annotated with BAKTA(v1.6.1) ^20^. Panaroo ^21^ was then used to define core and accessory genes. A core gene was defined as being present in 99% of all sequenced isolates.

Using the presence/absence gene matrix generated by Panaroo, we calculated Jaccard distances between accessory genomes of all pairs of isolates as:

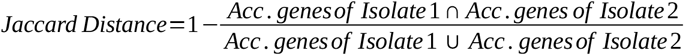

Pangenome accumulation curves were obtained using the R micropan package ^22^.

### Population structure determination

Multi-locus sequence type of isolates was predicted *in silico* with mlst (v2.1.6) (https://github.com/tseemann/mlst). Phylogroups were predicted with ClermonTyper (v20.03) (https://github.com/A-BN/ClermonTyping). PopPUNK (v2.4) ^23^ was used to assign draft genomes to existing clusters according to the *E. coli* database (v1) available at (https://www.bacpop.org/poppunk/).

A Neighbour-joining tree was constructed using IQ-TREE (v2.2.0.3), based on a core-genome alignment obtained with Panaroo.

### Plasmidome analysis and plasmid reconstructions

The plasmidome of each genome was defined as all plasmid-predicted contigs identified by using plasmidEC (v1.3) ^24^. Similar to previously described, plasmidomes were annotated with BAKTA, and Jaccard distances between these were calculated based on the presence/absence gene matrix generated by Panaroo.

Individual plasmids were reconstructed using gplas2 (v1.0) ^24^. Distances between all plasmid predictions were obtained using MASH (v2.2.2) ^25^ with k-mer length of 21, and a sketch size of 10,000. Clusters of highly similar plasmids were obtained by creating a network in which connections between plasmids were drawn if their MASH distance was below 0.01.

Clusters of plasmids backbones were created using mge-cluster(v1.1) ^26^ and clusters numbers were assigned based on the existing *E. coli* database, which can be accessed at: https://doi.org/10.6084/m9.figshare.21674078.v1

### Comparison of plasmidome and accessory genomes

Distances between accessory genomes and plasmidomes of isolates were visualized using t-distributed stochastic neighbor embedding algorithm (t-SNE), as implemented in the Rtsne R package (v0.15).

To conduct permutational analysis of variance (PERMANOVA), we used the adonis function from the vegan R package (v2.5-6) using the matrix of pairwise Jaccard distances as input. To explain the variance of accessory genome and plasmidome distances, six different PERMANOVA models were built with different explanatory variables each, as detailed in Supplementary Tables S1 and S2. In models with two variables, interaction terms between them are indicated with ‘*’.

### Estimation of plasmid copy number

After short-read assembly with unicycler, each contig is assigned a relative coverage value. We used all unitigs that unambiguously aligned to a single replicon to calculate the mean relative coverage of each plasmid. Duplicated contigs, aligning to more than one location of the genome, were left out of these calculations.

### ARGs, co-occurrence networks, and genomic context

Antibiotic resistance genes were identified by using AMRFinderPus (v3.11.2). Co-occurence of ARGs in the same plasmids were calculated by using a previously described approach ^27^. The genomic context of the tobramycin resistance transposon and of *bla*_CTX-M-15_ genes were obtained by manually exploring the assembly graph, and using BLAST against the ISFinder database of the nodes that surrounded the aforementioned elements.

### Statistics and code availability

Comparison of medians was performed using the non-parametric test Wilcoxon rank sum test ^28^ and Fisher’s exact test ^29^, respectively. All statistical analysis was performed using R (v3.6.1) and code needed to reproduce this analysis can be found in: https://gitlab.com/jpaganini/rgnosis_sdd_baseline.

## Results

### Patients colonised with ESBL *E. coli* in five European ICUs

The five selected ICUs (AN, PS, UZ, LB and CD) were located in five different hospitals in Belgium (n=3), Spain (n=1) and the UK (n=1). In total, 129 isolates were obtained from 116 patients, and in most cases (n=103, 90.6%) a single isolate per patient was sequenced. Most sequenced isolates derived from the ICU termed LB (n=55), while the rest of isolates were similarly distributed across the remaining locations ranging from 16 (PS) to 21 (AN) isolates per ICU (Table 1). There was a similar number of isolates from SDD (n=63) and baseline (n=66) periods. The majority of isolates (n=124, 96.1%) were obtained from rectal swabs, while a small number derived from respiratory samples (n=4, 3.1%) and bloodstream infections (n=1, 0.8%). Also, the majority of isolates (n=122, 94.6%) were phenotypically resistant to cefotaxime, ceftriaxone and/or ceftazidime. We refer to the isolates as ESBL-*E. coli, a*lthough not all isolates have phenotypic ESBL confirmation. The median time between the start of SDD and sample collection was 4 days (IQR=2–6.5), while for baseline samples, the median time was 6 days (IQR=4–11) after inclusion. All metadata associated with patients and sequenced isolates can be found in Supplementary Data 1.

**Table 1.**
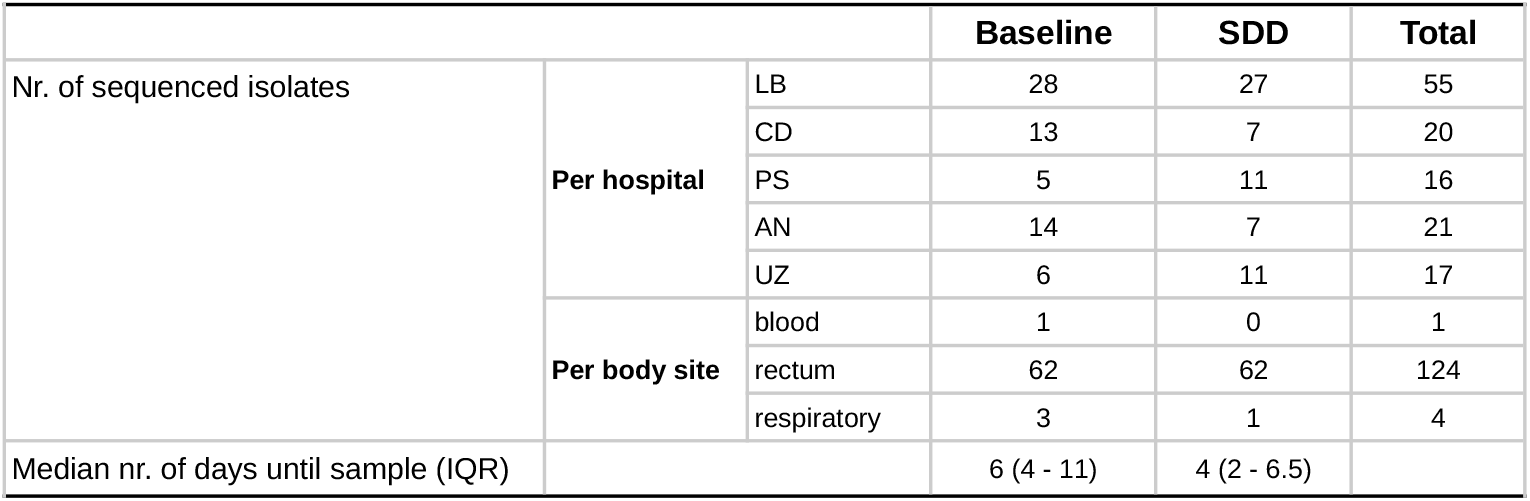
Sequenced isolates according to study period, hospital and time of isolation.

|  |  |  | Baseline | SDD | Total |
| --- | --- | --- | --- | --- | --- |
| Nr. of sequenced isolates | Per hospital | LB | 28 | 27 | 55 |
|  |  | CD | 13 | 7 | 20 |
|  |  | PS | 5 | 11 | 16 |
|  |  | AN | 14 | 7 | 21 |
|  |  | UZ | 6 | 11 | 17 |
|  | Per body site | blood | 1 | 0 | 1 |
|  |  | rectum | 62 | 62 | 124 |
|  |  | respiratory | 3 | 1 | 4 |
| Median nr. of days until sample (IQR) |  |  | 6 (4 - 11) | 4 (2 - 6.5) |  |

### Population structure of colonising ESBL-*E. coli*

Sequenced isolates belonged to 54 different STs, 11 of these were present in both study periods, 24 STs were only found in isolates from baseline period and 19 were exclusively found in SDD patients (Supplementary data 1). ST131 was the predominant clone (n=30) in both groups (baseline n=16, 24%; SDD n=14, 22%) (Figure 1A), followed by ST410 (n=7), ST10 (n=6) and ST1193 (n=5). We used PopPUNK to assign isolates to existing clusters considering both core and accessory genome variations (Supplementary data 1). We found a total of 53 clusters, of which the most abundant was Cluster 2 (n=27, 21%), which was entirely composed of ST131 isolates. Cluster 7_510, the second most abundant (n=12, 9.3%), encompasses isolates from ST10 (n=4), ST167 (n=3), ST744 (n=4) and ST1695 (n=1). A more detailed exploration of the core-genome of isolates (Figure 1B) showed no clear clusters associated with treatment, suggesting that SDD does not select for particular *E. coli* clones or lineages.

**Figure 1.**
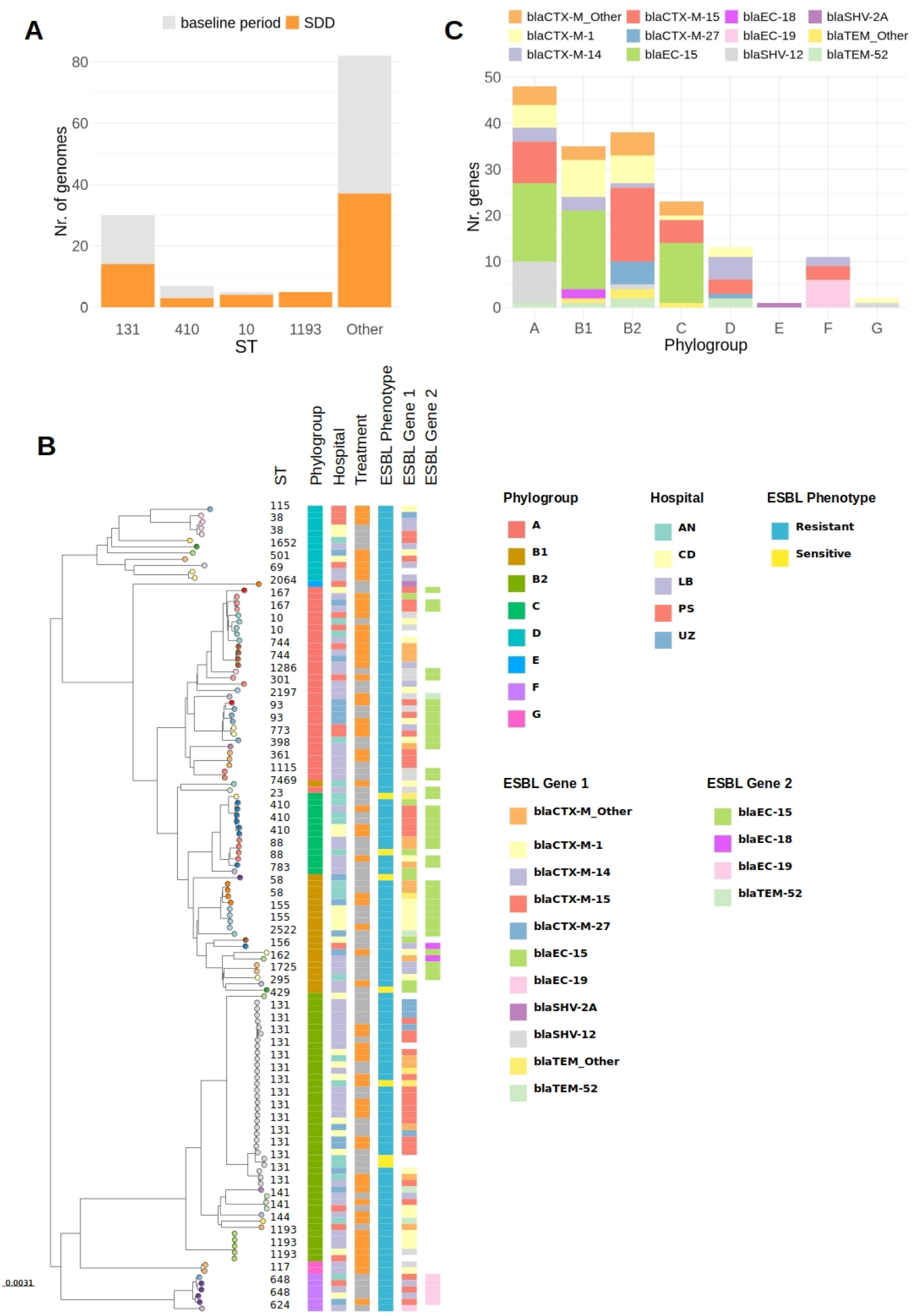
**A)** Distribution of most abundant ST across study periods. Groups with less than 5 isolates were collapsed into the ‘Other’ category. **B)** Neighbor-joining phylogenetic tree constructed based on core-genome alignment. Labels on the leaves indicate ST of isolates. Phylogroups were predicted *in silico* using ClermonTyper. ESBL Genes were predicted with AMRFinderPlus, if a second ESBL gene was present in an isolate, this is indicated in the column ESBL_Gene_2. ESBL phenotype indicates phenotyipic resistance or sensitivity to third-generation cephalosporins, evaluated as indicated in Methods. **C)** Distribution of the different ESBL genes across phylogroups.

Simpson’s indices of diversity calculated using ST (baseline=0.919, SDD=0.921) and PopPUNK clusters (baseline=0.924, SDD=0.918) showed that the population structure in both study periods was equally diverse.

Predictions of ARGs from our dataset showed that most isolates (n=123, 95.3%) carried at least one ESBL gene (Figure 1B). All isolates classified as phylogroup B2 (n=42) carried only one ESBL gene, with *bla*_CTX-M-15_ being the most abundant allele (n=16, 38.1%), followed by *bla*_CTX-M-1_ (n=6, 14.3%) and *bla*_CTX-M-27_ (n=5, 11.9%) (Figure 1B and C). Similarly, samples belonging to phylogroup D (n=12) also carried one ESBL gene, but *bla*_CTX-M-14_ was the most prevalent variant (n=5, 41.6%). In contrast, isolates from phylogroups A (n=17/30), B1 (n=15/20) and C (n=10/13) carried two ESBL genes, and *bla*_EC-15_ was the most frequent gene in all phylogroup, with prevalences of 35%, 48% and 57%, respectively.

### Variations in accessory genome compositions are associated with phylogroup and hospital, but not with the use of SDD

Considering all 129 isolates, the total pangenome consisted of 13,753 genes, of which 3,072 were identified as core- and the remaining 10,681 as accessory-genes. Accumulation curves fitted to Heap’s law for baseline and SDD isolates yielded similar alpha values (baseline=0.87, SDD=0.91) (Supplementary Figure S1A), confirming an open pangenome for both groups ^30^. The number of accessory genes in isolates from the baseline (median=1,645, IQR=1,503 - 1,776) and SDD periods (median=1,642, IQR=1,476-1,775) did not differ (p-value=0.92, Wilcoxon rank sum test) (Supplementary Figure S1B).

To identify associations between gene frequency and function, we obtained the functional categories of Clusters of Orthologous Groups (COG) from BAKTA annotations. A COG function was assigned to 66.1% of all predicted coding sequences (CDS). We performed a Fisher’s exact test to compare the frequency of each COG category in baseline and SDD isolates. This analysis showed that the distribution of genes to COG categories was similar between isolates from both study periods (Supplementary Figure S1C, Supplementary Table S3).

Next, we explored the diversity in total accessory gene content of individual isolates and its association with phylogroup, hospital and treatment by using PERMANOVA (Figure 2A and 2B, Supplementary Table S1). The accessory genome composition was strongly associated with the isolate’s phylogroup (R2=0.439, p-value=0.001), and its interaction with the geographical location (R2=0.10, p-value=0.006). There was no significant effect of SDD in the accessory genome composition (R2=0.01, p-value=0.149).

**Figure 2.**
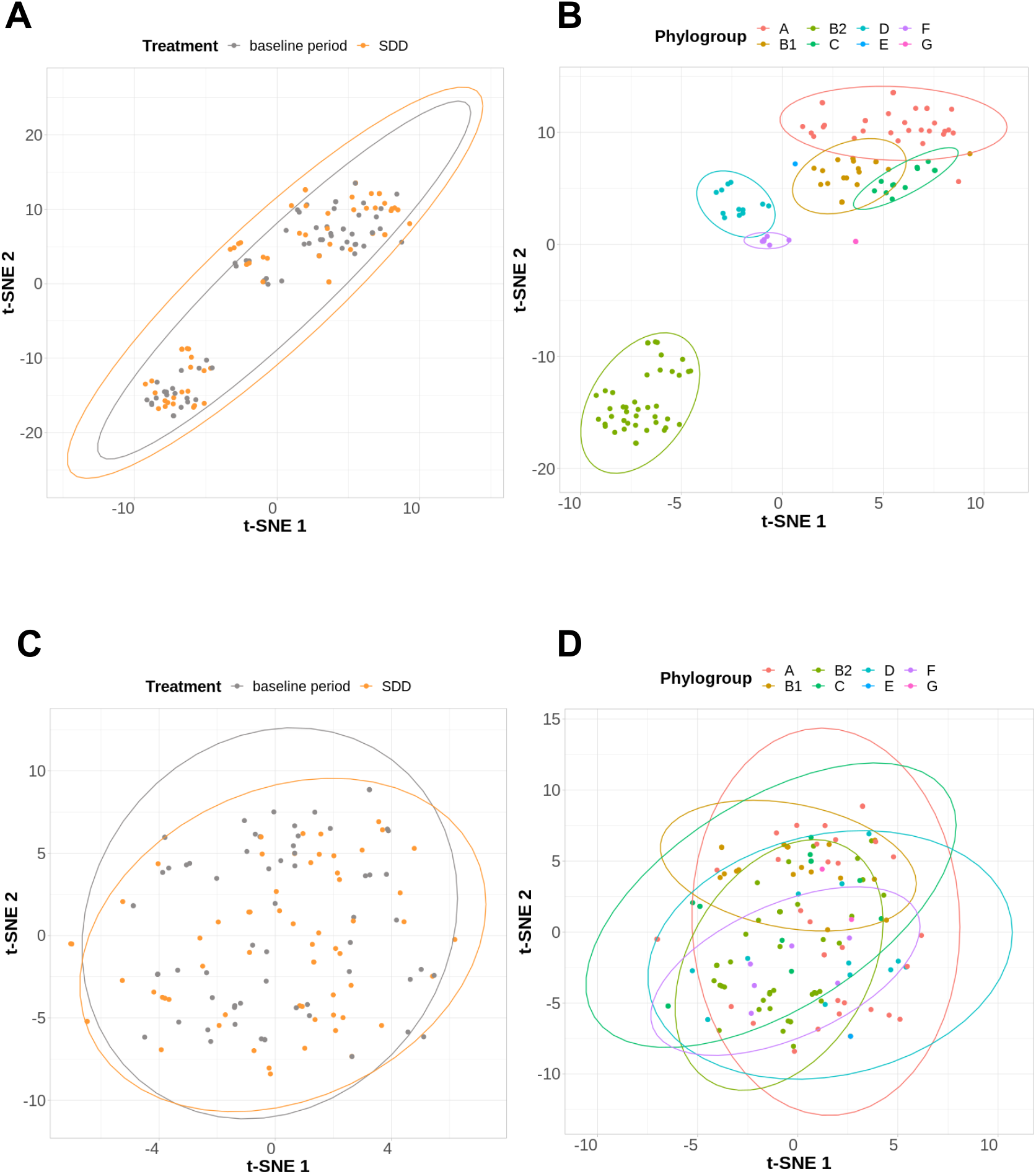
t-SNE plots representing Jaccard distances between complete accessory genomes **(A and B)** and predicted plasmidomes **(C and D)** for 129 ESBL-*E. coli* genomes included in this study. Jaccard distances were calculated using gene presence/absence.

The plasmidome is important for niche adaptation in *E. coli* ^31^ and other gut bacteria ^32^. Consequently, we also compared predicted plasmidome compositions across study periods. The median of plasmidome sizes for baseline isolates was 196,520.5 bp (IQR= 144,476.5 - 277,834.5) and of 216,196.0 bp (IQR= 141,770.0 – 293,638.8) for SDD isolates, which was not statistically different (p-value=0.92, Wilcoxon rank sum test, Supplementary Figure S2A). Additionally, the median number of unique plasmids per isolate, as predicted by gplas2 reconstructions, was 4 in isolates from both study periods (p-value=0.92, Wilcoxon rank sum test, Supplementary Figure S2B). Sizes and copy number of individual plasmids also followed expected distributions in both study periods ^33^ (Supplementary Figure S2C).

The use of SDD was also not associated with plasmidome variation (R2=0.011, p-value=0.075, PERMANOVA), but the interaction between phylogroup and the ICU explained the largest fraction of plasmidome variation (R2=0.154, p-value=0.002, PERMANOVA) (Figure 2C and 2D, Supplementary Table S2).

Since the ICU had a significant effect on the plasmidome composition, we wanted to evaluate if this was related to the fact that highly specific plasmid (or clones) were persistent over time in each ICU. For this, we predicted individual plasmids using gplas2, and clustered these plasmid predictions based on MASH distances (Supplementary Figure S3A and S3B). A total of 558 plasmids were predicted in 128 isolates. Of these, 257 plasmids (46%) were grouped into 65 clusters composed of highly similar plasmids (MASH distance < 0.01), and 37% of these clusters were exclusively composed of plasmids isolated from a single hospital, while 63% included plasmids from multiple hospitals. Moreover, 40% (n=12) of plasmid clusters that were recovered in single hospitals were found in multiple clones (PopPUNK clusters) (Supplementary Figure S3C, Supplementary data 3).

### A putative mobile genomic element encoding a tobramycin resistance gene is enriched in isolates from SDD treated patients

A total of 100 unique ARGs were found in the entire dataset (Supplementary data 4). Isolates obtained during baseline treatment contained a similar number of acquired ARGs (median=12, IQR = 8 - 14) as those of SDD (median=11, IQR= 7 - 13) (p-value=0.27, Wilcoxon ranked-sum test) (Supplementary figure S4). When subclassifying ARG by antibiotic class, we found a higher number of aminoglycoside resistance genes in baseline isolates (median= 3, IQR=1.25 - 4) than in SDD isolates (median=2, IQR= 1 - 3) (p-value=0.03, Wilcoxon ranked-sum test). For other ARG classes, no significant differences across study periods were found.

We then compared the occurrence of individual ARGs (Supplementary Figure S5) in both study periods. A total of 12 ARGs had an absolute difference in prevalence larger than 10% across study periods. Out of these, six genes were more frequent in baseline isolates, including two that code for beta-lactamases - *bla*_EC-15_, *bla*_TEM-1_ -, two genes that provide resistance to streptomycin [*aph(6)-ld, aph(3”)-lb*], and two ARGs that provide resistance to sulfonamide (*sul2*) and tetracyclin (*tet(A)*). Additionally, two genes were more frequent during SDD, namely *aac(6’)-Ib-cr5* and *qnrS1*.

The *aac(6’)-Ib-cr5* encodes an aminoglycoside 6’-N-acetyltransferase, which is predicted to confer resistance to tobramycin, one of the components of SDD ^1^. This gene was detected in 5/66 (7.5%) isolates during baseline and in 14/63 (22.2%) isolates during SDD. For other tobramycin-resistance genes, the absolute differences in prevalence between the baseline and SDD periods were below 5% (Figure 3A). In the majority of SDD isolates (n=13/14,92.9%), *aac(6’)-Ib-cr5* was encoded in conjunction with *bla*_OXA-1_ and with *catB3* on a ∼2.2Kb contig (Figure 3B), which was flanked by IS26 elements (Supplementary Figure S6). This suggests that this element is a potentially mobile composite transposon, termed from here on Tn(TobraR).

**Figure 3.**
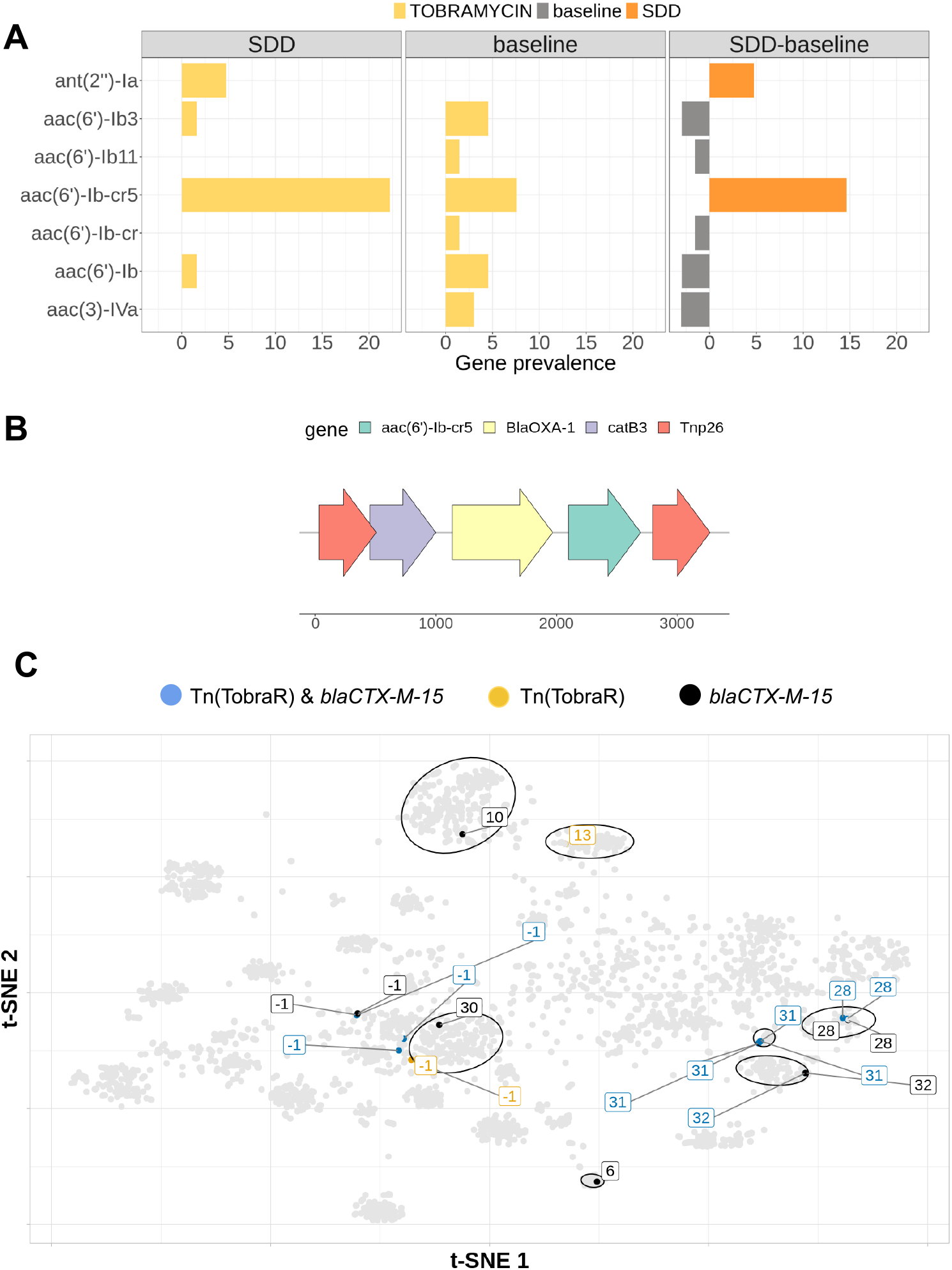
**A)** Prevalence of 7 ARGs predicted to provide resistance against tobramycin, one of the components of SDD treatment, in isolates from baseline (left) and SDD (center) patients. The difference in prevalence between the two study periods is displayed on the most right panel. **B)** Putative transposon carrying the *aac(6’)-Ib-cr5* gene, termed Tn(TobraR). **C)** t-SNE plot in which each dot represents an *E. coli* plasmid. Grey dots represent complete plasmids obtained from NCBI database (n∼4,500). Coloured dots represent plasmid predictions of isolates from patients that received SDD treatment and that carry Tn(TobraR), *bla*_CTX-M-15_ or both. Labels and ellipses depict different plasmid types, obtained with mge-cluster (v1.1). Labels equal to [-1] correspond to plasmid not assigned to any plasmid type.

To evaluate if Tn(TobraR) was found in different plasmid backbones, we used mge-cluster to assign the plasmid predictions generated by gplas2 to existing *E. coli* plasmid clusters (see methods). We found Tn(TobraR) in four distinct plasmid backbones (Figure 3C), namely cluster 13 (n=1), 28 (n=2), 31 (n=4) and 32 (n=1), and also in 5 different plasmids that were not assigned to a previously existing plasmid type, supporting the hypothesis that Tn(TobraR) can actually move independently. Additionally, this element was found in isolates with different chromosomal backgrounds, including phylogroups B2 (n=9, most ST131), C (n=1), F (n=1) and A (n=2) (Supplementary Figure S7A). Moreover, when querying a database composed of more than 1,300 publicly available *E. coli* genomes, we found the Tn(TobraR) in 63 additional genomes from six different phylogroups (Supplementary Figure S7B). Notably, in SDD isolates, we observed that the ARGs that compose the Tn(TobraR) co-occurred with *bla*_CTX-M-15_ in the same plasmid significantly more frequently than expected by chance (Supplementary Figure S8A, Supplementary table S4). However, this was not the case in baseline isolates (Supplementary Figure S8B, Supplementary table S5).

Finally, we evaluated the prevalence of genes coding for carbapenem resistance. A total of four isolates were predicted to have a carbapenemase gene, all of which were obtained from SDD-treated patients. One of these isolates carried a *bla*_OXA-48_ gene in an ST295 background; two isolates from the same patient, belonging to ST410, coded a *bla*_OXA-181_ gene; and *bla*_VIM-1_ was found in an ST1193 isolate.

## Discussion

In this work we sequenced the genomes of 129 ESBL-*E. coli* isolates of which 63 were obtained from ICU patients that received SDD as a prophylactic measure to prevent colonisation and infection with potentially pathogenic gram-negative bacteria, and compared those with 66 genomes of ESBL-*E. coli* isolates from the same patient population that did not receive SDD. One of the strengths of our study is its multi-center nature, which ensures broader representativeness by including isolates from five different ICUs located in three European countries, reducing bias associated with single-center studies. However, the multi-center nature also introduces logistical complexity, as differences in sample handling or metadata collection across centers that could introduce unexpected variability. Moreover, in contrast to many similar studies ^34–38^, 96% of the isolates represented intestinal carriage isolates, rather than being recovered from clinical samples. This design avoids clinical bias and offers valuable insight into the genetic and resistome features of *E. coli* isolates colonizing the gut. Nonetheless, a limitation of this approach is the lack of data on unaccounted temporal changes, as isolates were collected at specific time points rather than longitudinally, potentially missing transient or early-colonizing strains.

The results from our study revealed no important differences in pangenome compositions between ESBL-*E. coli* recovered from patients treated with SDD and those not treated with SDD. These findings do not support a large impact of SDD in shaping the overall pangenome composition of ESBL-*E. coli* after a median of four days of SDD exposure. These results were unexpected when considering that SDD alters the microbial composition of the gut ^17,18^, potentially changing the interaction networks that occur in the microbiota leading to new metabolic challenges for *E. coli* ^39^. The absence of important differences in the pangenome can have several potential explanations. First, it is possible that the adaptation of ESBL-*E. coli* to the new gut ecology induced by SDD is not mediated by the acquisition/loss of certain genes, but rather by changes in gene expression patterns. These changes cannot be detected by our analysis which solely relies on gene content comparisons. Supporting this hypothesis, a recent study has described that a global re-wiring of transcription factors is observed when switching *E. coli* from auxotrophic to prototrophic growing conditions ^40^. Moreover, another study demonstrated that *E. coli* auxotrophies can be rescued by expressing short peptides that are coded in novel small open reading frames ^41^, which will also be missed by the annotation tools that we have used in this study. Second, it is also possible that the duration of SDD was not sufficient to cause an impact in the community structure of *E. coli*. Finally, the fact that we had only ESBL-positive isolates available for WGS, prohibited analysis of changes in the relative abundances of different *E. coli* subpopulations within each patient. Future research should ideally collect multiple isolates (or feacal samples) over longer periods of time from the same patients, including also non-ESBL *E. coli*.

The interplay between phylogeny and the ICU explained the largest fraction of variance observed in the accessory genome and plasmidome of isolates. The strong association between phylogeny and the accessory genome of *E. coli* has already been described in ^31,42^. The effect of the ICU could be explained by postulating that each ward constitutes its own ecosystem, in which particular plasmids, and clones, persist over time, with the ability to spread to different patients. In line with this hypothesis, recent studies suggest that plasmids carrying carbapenem resistance genes present ‘geographical signatures’ that relate them to particular healthcare settings ^43^. Moreover, the long-term persistence of clones and plasmids in clinical environments is more common than originally thought ^44–48^. A patient admitted to an ICU can be colonised by bacteria contaminating this environment in less than 6 days ^44^. On the broader scale, particular plasmids have also been associated with specific countries ^49,50^.

We also observed minor differences between the resistomes of SDD and baseline isolates. While six different resistance genes were more prevalent in baseline isolates, SDD isolates were enriched for a potentially mobile genetic element composed of a tobramycin resistance gene and two additional ARGs, flanked by IS26 elements. An identical genetic element was reported in other studies ^51,52^, but it mobility as an independent unit was never tested. It is well known that IS26 plays a crucial role in the dissemination of resistance genes among Enterobacteriaceae in the clinical environment^53–55^. IS26 catalyses a highly efficient conservative transposition reaction that allows the incorporation of ARGs preferably into replicons that contain a pre-existing copy of this element ^56,57^. This mechanism could lead to the formation of arrays of in-tandem resistance genes, also referred to as resistance islands ^53^. In line with this, we observed that the tobramycin resistant transposon identified in this work frequently co-occurred with a *bla*_CTX-M-15_ gene in multiple plasmid backbones that reside in distinct *E. coli* clones. This means that if SDD in fact selects for this transposon, it could also facilitate the formation of resistance islands that accommodate multiple ARGs. However, it is important to note that we had insufficient isolates to perform multivariable analysis that can correct for population structure, such as a bacterial GWAS ^58^. Moreover, it should be noted that we have not collected data on the types and amount of therapeutic antibiotics used in these patients, so differences in the resistome should be interpreted with care.

A previous study based on the R-GNOSIS ICU data found that phenotypic resistance to colistin was rare ^13^, despite this being one of the antibiotics administered in SDD. In concordance with this result, no mcr genes were predicted based on WGS data, and only four isolates with phenotypic resistance were reported. Nevertheless, phenotypic resistance was determined using the E-test method, which has limited predictive accuracy according to Galani et al ^59^.

Overall, our study constitutes the first WGS-based analysis of the potential effects of SDD in the pangenome composition of a PPMO. Despite the limitations in sample collection, our results suggest that SDD treatment has limited effects in the accessory genome, plasmidome and resistome compositions of ESBL-*E. coli*.

## Notes

### Competing Interest Statement

The authors have declared no competing interest.

