## Supplementary Material for "Impact of selective digestive decontamination on the pangenome composition of ESBL-*E. coli*"

### Supplementary Materials

#### Supplementary Data

Supplementary data can be downloaded from: https://doi.org/10.5281/zenodo.7926667

#### Supplementary Tables

**Supplementary Table S1.** Results of PERMANOVA analysis to model the variance observed in accessory genome composition. The ‘*’ symbol indicates interaction term between variables. Treatment codes for SDD vs baseline.

| Model nr. | Explanatory variables | Df | F.Model | R2 | Pr(>F) |
|---|---|---|---|---|---|
| 1 | Phylogroup | 7 | 13.5281039 | 0.4390271864 | 0.001 |
| 2 | Hospital | 4 | 1.216606935 | 0.03776334787 | 0.154 |
| 3 | Treatment | 1 | 1.365392476 | 0.01063676471 | 0.149 |
| 4 | Phylogroup | 7 | 13.7654975 | 0.4390271864 | 0.001 |
| | Treatment | 1 | 1.484859951 | 0.00676529839 | 0.102 |
| | Phylogroup*Treatment | 5 | 1.327693956 | 0.03024610429 | 0.054 |
| 5 | Phylogroup | 7 | 14.31769118 | 0.4390271864 | 0.001 |
| | Hospital | 4 | 1.430689111 | 0.02506834405 | 0.054 |
| | Phylogroup*Hospital | 18 | 1.296643498 | 0.102238266 | 0.006 |
| 6 | Treatment | 1 | 1.372662651 | 0.01063676471 | 0.143 |
| | Hospital | 4 | 1.2108664 | 0.03753202137 | 0.148 |
| | Treatment*Hospital | 4 | 0.9581897992 | 0.0297000561 | 0.532 |

**Supplementary Table S2.** Results of PERMANOVA analysis to model the variance observed in plasmidome composition. The ‘*’ symbol indicates interaction term between variables. Treatment codes for SDD vs baseline.

| Model nr. | Explanatory variables | Df | F.Model | R2 | Pr(>F) |
|---|---|---|---|---|---|
| 1 | Phylogroup | 7 | 2.262016978 | 0.115717622 | 0.001 |
| 2 | Hospital | 4 | 1.182226021 | 0.03673537126 | 0.099 |
| 3 | Treatment | 1 | 1.413281789 | 0.01100572908 | 0.072 |
| 4 | Phylogroup | 7 | 2.311171098 | 0.115717622 | 0.001 |
| | Treatment | 1 | 1.346775064 | 0.009633063636 | 0.083 |
| | Phylogroup*Treatment | 5 | 1.456516246 | 0.05209004107 | 0.003 |
| 5 | Phylogroup | 7 | 2.362907434 | 0.115717622 | 0.001 |
| | Hospital | 4 | 1.346915624 | 0.03769251651 | 0.016 |
| | Phylogroup*Hospital | 18 | 1.222732122 | 0.1539779659 | 0.002 |
| 6 | Treatment | 4 | 1.18561335 | 0.03673537126 | 0.085 |
| | Hospital | 1 | 1.528465622 | 0.01183960017 | 0.042 |
| | Treatment*Hospital | 4 | 0.9567051965 | 0.02964281785 | 0.592 |

**Supplementary Table S3.** Fisher's test results for abundance of different COG types across study periods.

| COG | p.value | Confidence interval | OR | Occurence SDD | Occurence baseline period | Not occurence SDD | Not occurence baseline period |
|---|---|---|---|---|---|---|---|
| J | 0.9852496345 | c(0.975789492902574, 1.02425758983737) | 0.9997054997 | 13722 | 14429 | 200445 | 210716 |
| K | 0.6056192133 | c(0.972489422695379, 1.0163625709264) | 0.9941842403 | 16759 | 17713 | 197408 | 207432 |
| F | 0.9719039845 | c(0.965246095111443, 1.03455419087637) | 0.9993147717 | 6486 | 6823 | 207681 | 218322 |
| Q | 0.9366409714 | c(0.951155658340286, 1.05601402039197) | 1.002227294 | 2819 | 2957 | 211348 | 222188 |
| G | 0.6018505376 | c(0.985794356531915, 1.02508083454845) | 1.005237299 | 22074 | 23097 | 192093 | 202048 |
| I | 0.6933838823 | c(0.974037533976659, 1.0405276915715) | 1.006727248 | 7196 | 7516 | 206971 | 217629 |
| E | 0.8130938607 | c(0.982364983479942, 1.02289454010526) | 1.00244109 | 20432 | 21432 | 193735 | 203713 |
| C | 0.9634520666 | c(0.978245285628295, 1.02330688097101) | 1.000524396 | 16079 | 16895 | 198088 | 208250 |
| M | 0.8085665345 | c(0.974179799777156, 1.02052782175703) | 0.9970788874 | 15003 | 15815 | 199164 | 209330 |
| O | 0.906893496 | c(0.969951310487423, 1.02736934293829) | 0.9982685969 | 9547 | 10053 | 204620 | 215092 |
| R | 0.8888010572 | c(0.972289137890958, 1.02457843874712) | 0.9980901027 | 11625 | 12243 | 202542 | 212902 |
| U | 0.8477019785 | c(0.952154529987101, 1.04089787909678) | 0.99555203 | 3884 | 4101 | 210283 | 221044 |
| T | 0.9945601485 | c(0.973350466549645, 1.02703215378781) | 0.9998562239 | 11065 | 11634 | 203102 | 213511 |
| P | 0.9345003041 | c(0.976456561365551, 1.026244226516) | 1.001042395 | 13000 | 13653 | 201167 | 211492 |
| H | 0.8402077964 | c(0.976601630259693, 1.02963648049424) | 1.002776834 | 11422 | 11976 | 202745 | 213169 |
| S | 0.9486169264 | c(0.96991901626744, 1.03340134536923) | 1.001155743 | 7806 | 8197 | 206361 | 216948 |
| D | 0.8987001219 | c(0.949242121483854, 1.06140674472677) | 1.003776748 | 2471 | 2588 | 211696 | 222557 |
| V | 0.9926284441 | c(0.963969778713874, 1.03672824250224) | 0.9996686522 | 5875 | 6178 | 208292 | 218967 |
| L | 0.5763402559 | c(0.960966481318246, 1.02237950636188) | 0.9912170141 | 8154 | 8645 | 206013 | 216500 |
| X | 0.0649580104 | c(0.996571842983943, 1.11875968712521) | 1.055920887 | 2362 | 2353 | 211805 | 222792 |
| N | 0.1746329295 | c(0.934173122750306, 1.01245725611287) | 0.9725566337 | 4723 | 5102 | 209444 | 220043 |
| W | 0.970997954 | c(0.928852378768879, 1.07256736623477) | 0.9981455043 | 1485 | 1564 | 212682 | 223581 |
| A | 0.6192491546 | c(0.740511742054748, 1.68314621513269) | 1.115649109 | 52 | 49 | 214115 | 225096 |
| Z | 1 | c(0.779773374511084, 1.29089404928233) | 1.003476705 | 126 | 132 | 214041 | 225013 |

**Supplementary Table S4.** Co-occurrence of ARGs in the same plasmid in SDD isolates. Co-occurrences with a p-value smaller than 0.01 were considered significant.

| ARG 1 | ARG 2 | ARG 1 Occurence | ARG 2 Occurence | Co-occurence | Prob. Co-occurence | Expected co-occurence | p-value |
|---|---|---|---|---|---|---|---|
| aac(3)-IId | blaTEM-1 | 7 | 21 | 4 | 0.014 | 1.4 | 0.03039 |
| aac(3)-IId | mph(A) | 7 | 24 | 4 | 0.016 | 1.6 | 0.04957 |
| aac(3)-IIe | aac(6')-Ib-cr5 | 11 | 13 | 4 | 0.013 | 1.4 | 0.0312 |
| aac(3)-IIe | blaCTX-M-15 | 11 | 18 | 8 | 0.019 | 1.9 | 2.00E-05 |
| aac(3)-IIe | blaOXA-1 | 11 | 14 | 4 | 0.015 | 1.5 | 0.04108 |
| aac(3)-IIe | catB3 | 11 | 14 | 4 | 0.015 | 1.5 | 0.04108 |
| aac(6')-Ib-cr5 | blaCTX-M-15 | 13 | 18 | 10 | 0.022 | 2.3 | 0 |
| aac(6')-Ib-cr5 | blaOXA-1 | 13 | 14 | 13 | 0.017 | 1.8 | 0 |
| aac(6')-Ib-cr5 | catB3 | 13 | 14 | 13 | 0.017 | 1.8 | 0 |
| aac(6')-Ib-cr5 | mph(A) | 13 | 24 | 7 | 0.029 | 3 | 0.01073 |
| aadA1 | aadA2 | 17 | 8 | 5 | 0.013 | 1.3 | 0.00286 |
| aadA1 | aadA5 | 17 | 25 | 0 | 0.04 | 4.1 | 1 |
| aadA1 | blaCTX-M-15 | 17 | 18 | 0 | 0.029 | 3 | 1 |
| aadA1 | dfrA17 | 17 | 31 | 0 | 0.05 | 5.1 | 1 |
| aadA1 | mph(A) | 17 | 24 | 0 | 0.038 | 4 | 1 |
| aadA5 | blaCTX-M-15 | 25 | 18 | 9 | 0.042 | 4.4 | 0.00842 |
| aadA5 | blaOXA-1 | 25 | 14 | 7 | 0.033 | 3.4 | 0.02319 |
| aadA5 | blaTEM-1 | 25 | 21 | 9 | 0.049 | 5.1 | 0.02964 |
| aadA5 | catB3 | 25 | 14 | 7 | 0.033 | 3.4 | 0.02319 |
| aadA5 | dfrA17 | 25 | 31 | 20 | 0.073 | 7.5 | 0 |
| aadA5 | mph(A) | 25 | 24 | 17 | 0.057 | 5.8 | 0 |
| aadA5 | sul1 | 25 | 33 | 21 | 0.078 | 8 | 0 |
| aph(3'')-Ib | aph(6)-Id | 19 | 19 | 19 | 0.034 | 3.5 | 0 |
| aph(3'')-Ib | blaTEM-1 | 19 | 21 | 10 | 0.038 | 3.9 | 0.00048 |
| aph(3'')-Ib | sul2 | 19 | 23 | 17 | 0.041 | 4.2 | 0 |
| aph(3'')-Ib | tet(A) | 19 | 20 | 10 | 0.036 | 3.7 | 0.00028 |
| aph(6)-Id | blaTEM-1 | 19 | 21 | 10 | 0.038 | 3.9 | 0.00048 |
| aph(6)-Id | sul2 | 19 | 23 | 17 | 0.041 | 4.2 | 0 |
| aph(6)-Id | tet(A) | 19 | 20 | 10 | 0.036 | 3.7 | 0.00028 |
| blaCTX-M-15 | blaOXA-1 | 18 | 14 | 11 | 0.024 | 2.4 | 0 |
| blaCTX-M-15 | catB3 | 18 | 14 | 11 | 0.024 | 2.4 | 0 |
| blaCTX-M-15 | dfrA17 | 18 | 31 | 9 | 0.053 | 5.4 | 0.04365 |
| blaCTX-M-15 | sul1 | 18 | 33 | 10 | 0.056 | 5.8 | 0.02113 |
| blaOXA-1 | catB3 | 14 | 14 | 14 | 0.018 | 1.9 | 0 |
| blaOXA-1 | mph(A) | 14 | 24 | 7 | 0.032 | 3.3 | 0.01802 |
| blaOXA-1 | sul1 | 14 | 33 | 8 | 0.044 | 4.5 | 0.03464 |
| blaTEM-1 | sul1 | 21 | 33 | 11 | 0.065 | 6.7 | 0.02609 |
| catB3 | mph(A) | 14 | 24 | 7 | 0.032 | 3.3 | 0.01802 |
| catB3 | sul1 | 14 | 33 | 8 | 0.044 | 4.5 | 0.03464 |
| dfrA17 | mph(A) | 31 | 24 | 12 | 0.07 | 7.2 | 0.01661 |
| dfrA17 | sul1 | 31 | 33 | 16 | 0.096 | 9.9 | 0.00573 |
| mph(A) | sul1 | 24 | 33 | 18 | 0.075 | 7.7 | 0 |
| sul2 | tet(A) | 23 | 20 | 9 | 0.043 | 4.5 | 0.01047 |

**Supplementary Table S5.** Co-occurrence of ARGs in the same plasmid in baseline isolates. Co-occurrences with a p-value smaller than 0.01 were considered significant.

| ARG 1 | ARG 2 | ARG 1 Occurence | ARG 2 Occurence | Co-occurence | Prob. co-occurence | Expected co-occurence | p-value |
|---|---|---|---|---|---|---|---|
| aadA1 | aadA2 | 14 | 10 | 4 | 0.011 | 1.2 | 0.02122 |
| aadA1 | aadA5 | 14 | 22 | 0 | 0.025 | 2.8 | 1 |
| aadA1 | dfrA17 | 14 | 29 | 0 | 0.032 | 3.6 | 1 |
| aadA2 | dfrA17 | 10 | 29 | 0 | 0.023 | 2.6 | 1 |
| aadA5 | dfrA17 | 22 | 29 | 11 | 0.051 | 5.7 | 0.00601 |
| aadA5 | mph(A) | 22 | 20 | 12 | 0.035 | 3.9 | 1.00E-05 |
| aadA5 | sul1 | 22 | 31 | 14 | 0.054 | 6.1 | 8.00E-05 |
| aadA5 | sul2 | 22 | 38 | 12 | 0.067 | 7.5 | 0.02302 |
| aph(3'')-Ib | aph(3')-Ia | 39 | 12 | 9 | 0.037 | 4.2 | 0.00339 |
| aph(3'')-Ib | aph(6)-Id | 39 | 39 | 39 | 0.121 | 13.6 | 0 |
| aph(3'')-Ib | blaTEM-1 | 39 | 41 | 20 | 0.127 | 14.3 | 0.01618 |
| aph(3'')-Ib | dfrA1 | 39 | 4 | 4 | 0.012 | 1.4 | 0.01324 |
| aph(3'')-Ib | dfrA17 | 39 | 29 | 3 | 0.09 | 10.1 | 0.99989 |
| aph(3'')-Ib | sul2 | 39 | 38 | 32 | 0.118 | 13.2 | 0 |
| aph(3')-Ia | aph(6)-Id | 12 | 39 | 9 | 0.037 | 4.2 | 0.00339 |
| aph(3')-Ia | blaTEM-1 | 12 | 41 | 9 | 0.039 | 4.4 | 0.00522 |
| aph(3')-Ia | sul2 | 12 | 38 | 9 | 0.036 | 4.1 | 0.0027 |
| aph(6)-Id | blaTEM-1 | 39 | 41 | 20 | 0.127 | 14.3 | 0.01618 |
| aph(6)-Id | dfrA1 | 39 | 4 | 4 | 0.012 | 1.4 | 0.01324 |
| aph(6)-Id | dfrA17 | 39 | 29 | 3 | 0.09 | 10.1 | 0.99989 |
| aph(6)-Id | sul2 | 39 | 38 | 32 | 0.118 | 13.2 | 0 |
| blaCTX-M-27 | sul1 | 4 | 31 | 4 | 0.01 | 1.1 | 0.00507 |
| blaTEM-1 | dfrA17 | 41 | 29 | 5 | 0.095 | 10.6 | 0.99784 |
| blaTEM-1 | sul2 | 41 | 38 | 21 | 0.124 | 13.9 | 0.00333 |
| blaTEM-1 | sul3 | 41 | 6 | 5 | 0.02 | 2.2 | 0.02412 |
| blaTEM-1 | tet(M) | 41 | 3 | 3 | 0.01 | 1.1 | 0.04677 |
| dfrA1 | sul2 | 4 | 38 | 4 | 0.012 | 1.4 | 0.01188 |
| dfrA17 | tet(A) | 29 | 35 | 4 | 0.081 | 9.1 | 0.9968 |
| mph(A) | sul1 | 20 | 31 | 15 | 0.049 | 5.5 | 0 |
| mph(A) | tet(A) | 20 | 35 | 10 | 0.056 | 6.2 | 0.04451 |
| tet(A) | tet(B) | 35 | 13 | 1 | 0.036 | 4.1 | 0.99459 |

#### Supplementary Figures

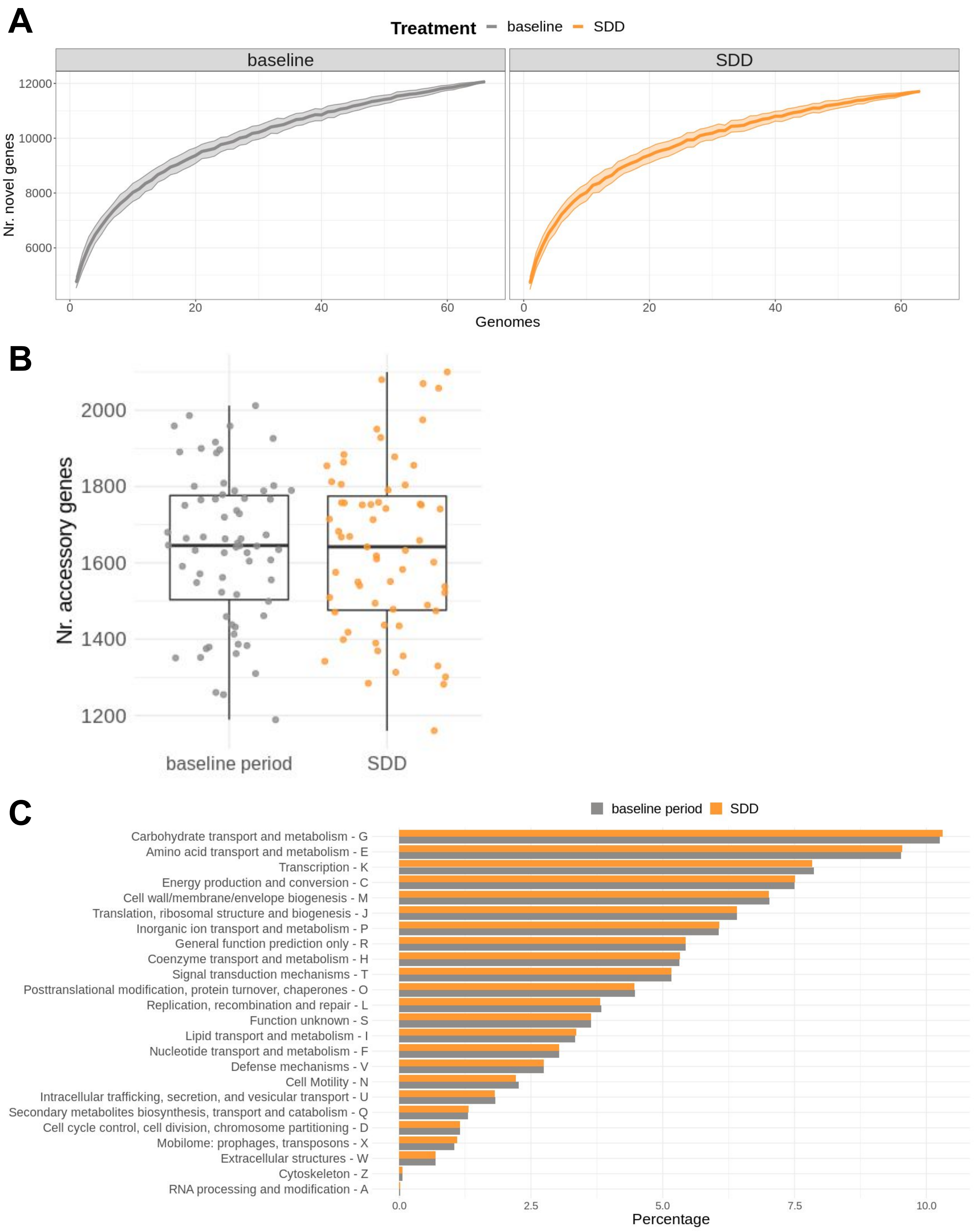


**Supplementary Figure S1. A)** Pangenome accumulation curves for baseline and SDD isolates. **B)** Number of accessory genes per isolate across different study periods. **C)** Fraction of COG functional categories by study period.

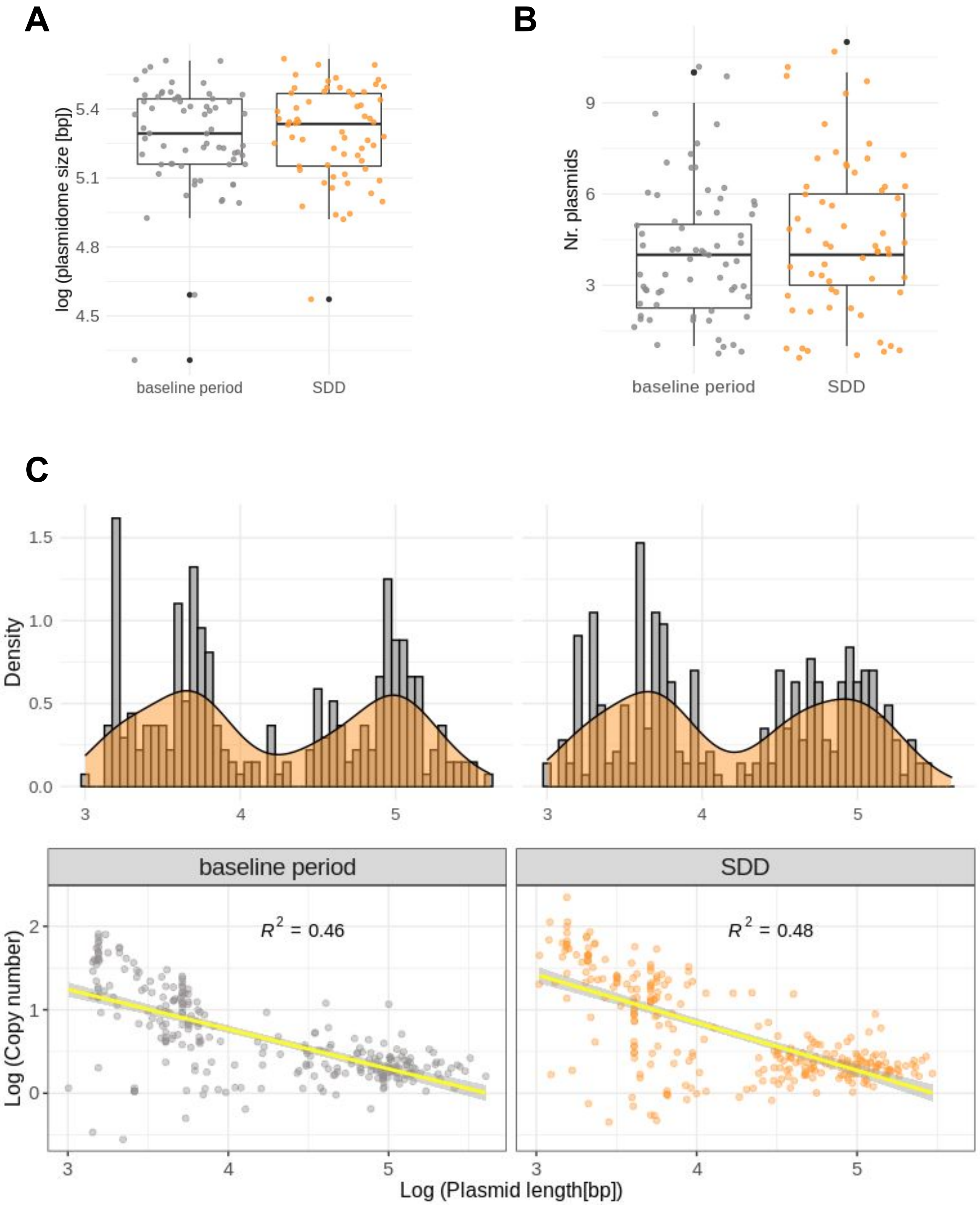


**Supplementary Figure S2. A)** Total size of predicted plasmidome sorted by study period. This is, the total length of all contigs within an isolate predicted to be plasmid by plasmidEC. **B)** Number of predicted individual plasmids per isolate. Plasmids were predicted using gplas. **C)** Plasmid size vs estimated copy number for individual plasmid predictions

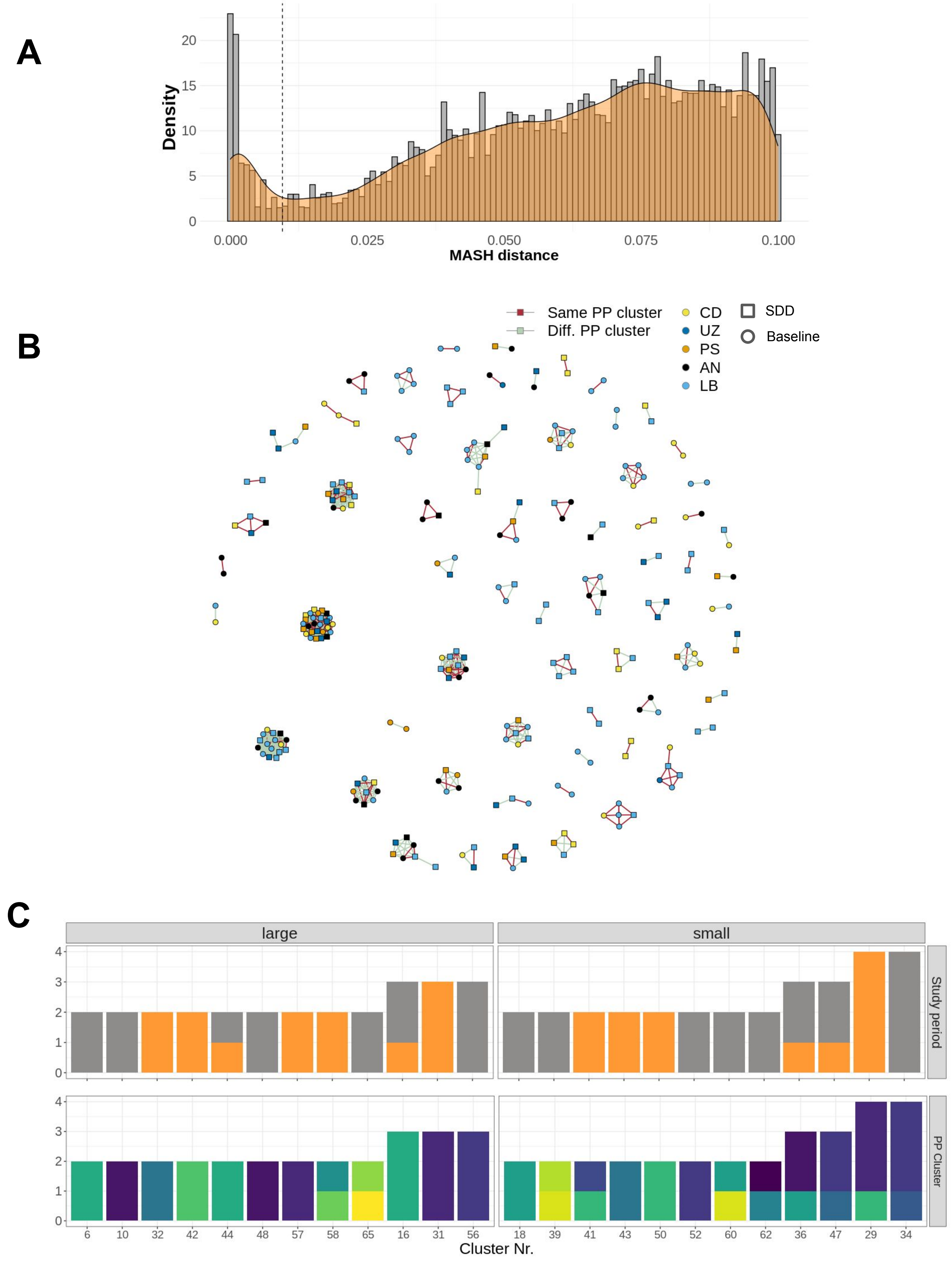


**Supplementary Figure S3. A)** MASH distances (k=21, s=10,000) for all plasmid-predictions vs plasmid-predictions. Dashed line indicates the cut-off point (distance =0.01) to create network of plasmids. **B)** Network displaying clusters of highly similar plasmids. **C)** For plasmid clusters that were present only in single hospitals, study period in which they were found and the number of distinct PopPUNK clusters is indicated.

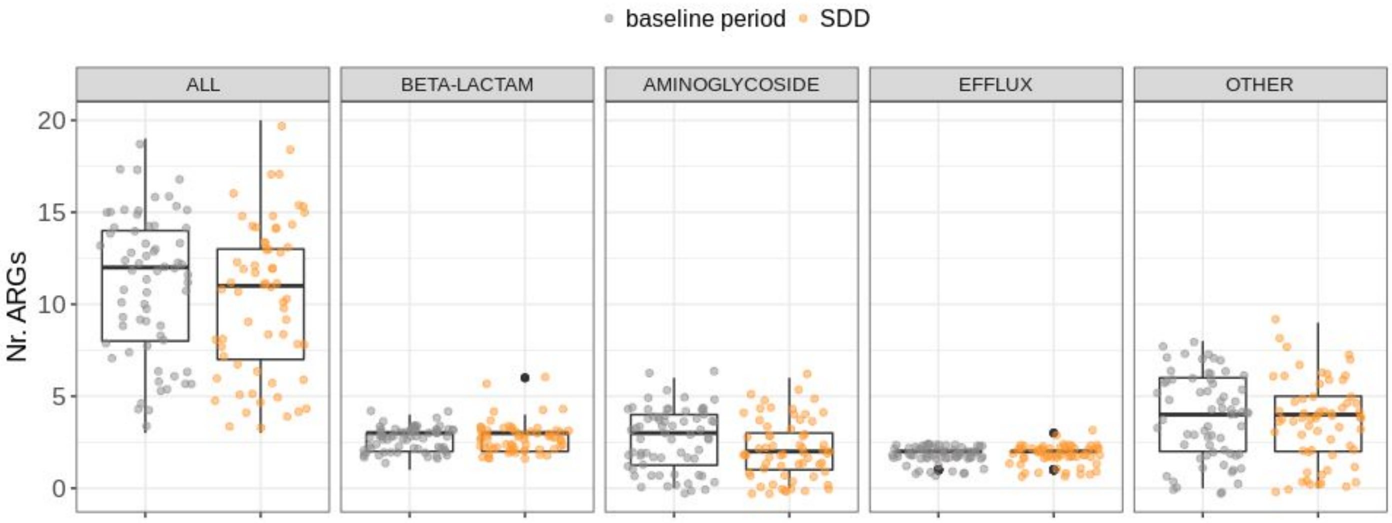


**Supplementary Figure S4.** Nr. of acquired ARGs per isolate, treatment and ARG type.

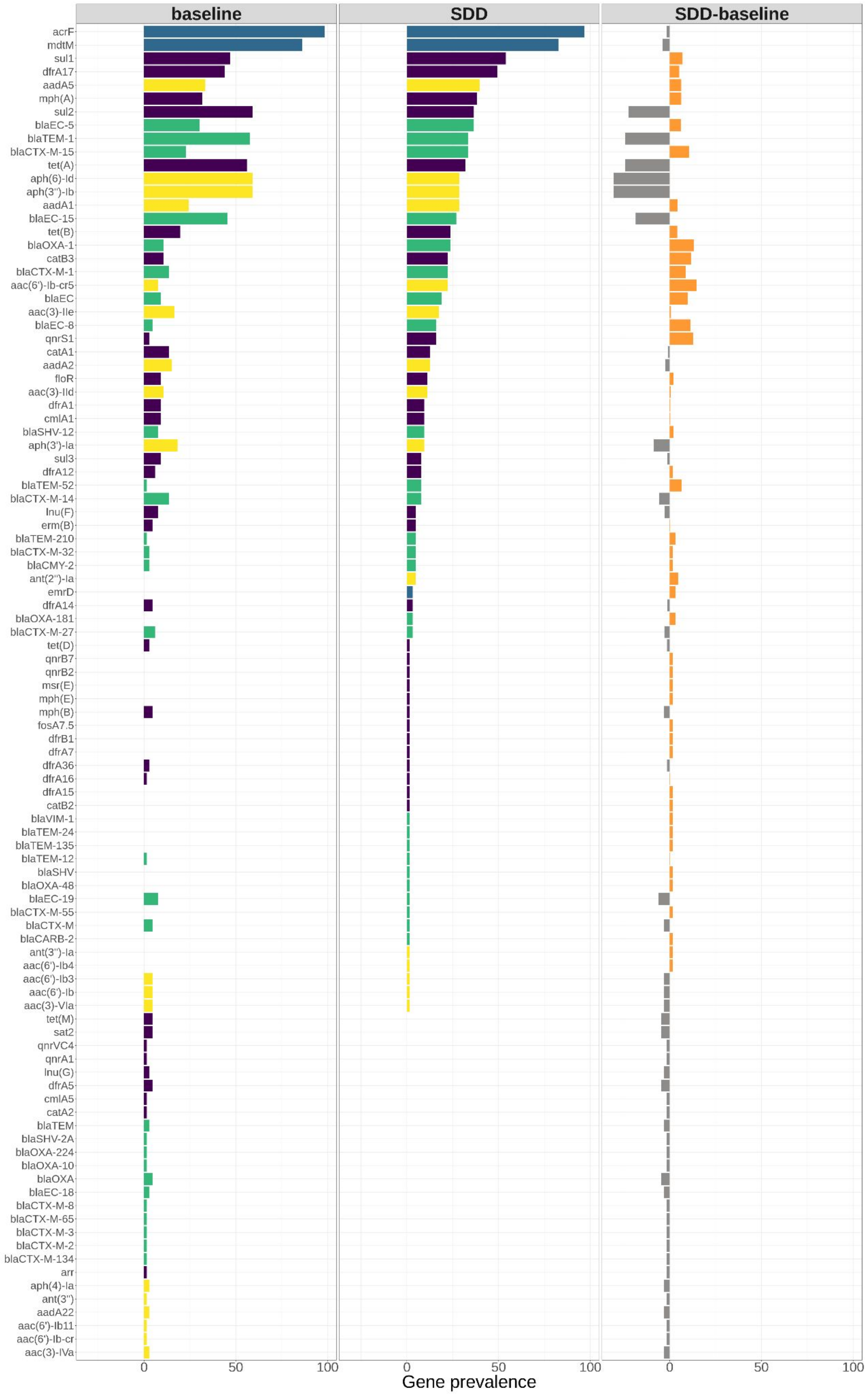


**Supplementary Figure S5.** The first two panels show the prevalence of all acquired ARGs in SDD and baseline isolates. The third panel shows the absolute difference between these prevalences in SDD and baseline periods.

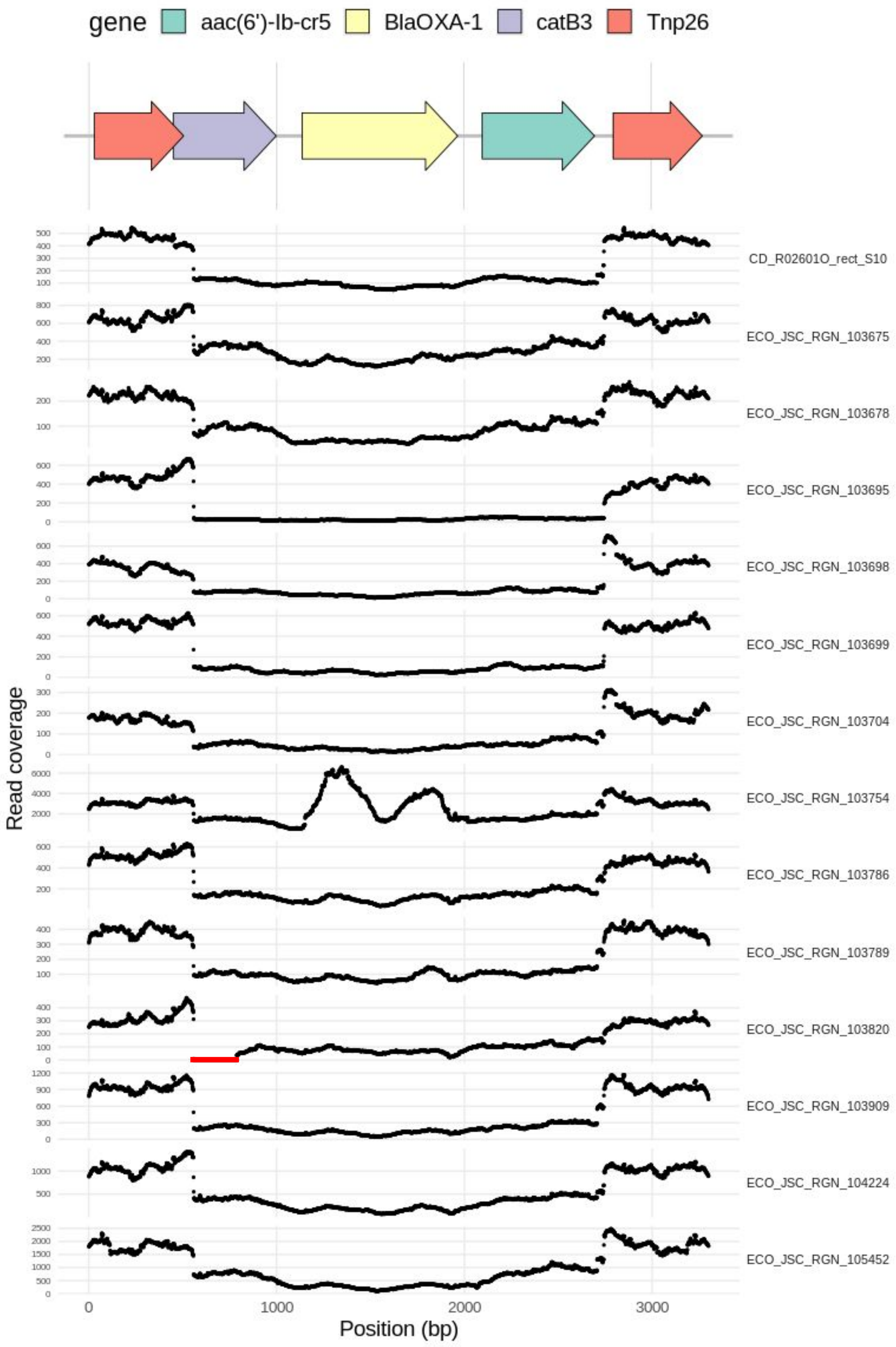


**Supplementary Figure S6.** Read coverage of each SDD isolate (n=14) that contains the putative transposon carrying the tobramycin resistance gene. Coverage suggests that all SDD isolates, except ECO-JSC-RGN-103820, carry the complete sequence of Tn(TobraR). Red lines indicate regiones with read coverage equal to zero.

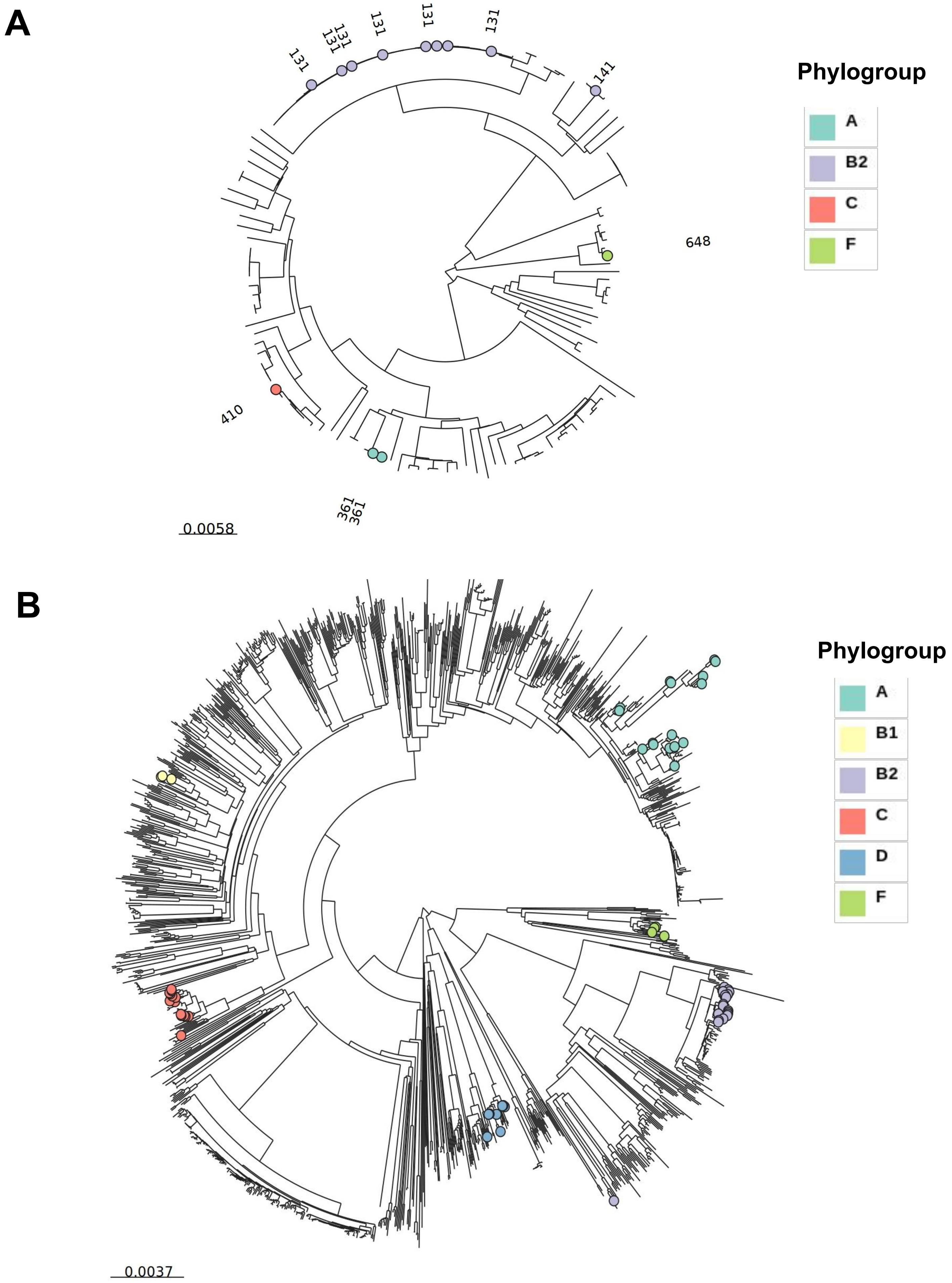


**Supplementary Figure S7. A)** NJ tree based on core-genome alignment of *E. coli* isolates from the R-GNOSIS study. Leaf labels indicate the sequence type of isolates. **B)** NJ cg-tree based on k-mer presence/absence of 1381 publicly available *E. coli* complete genomes. In both trees, colored nodes indicate the genomes that carry Tn(TobraR) and their corresponding phylogroup.

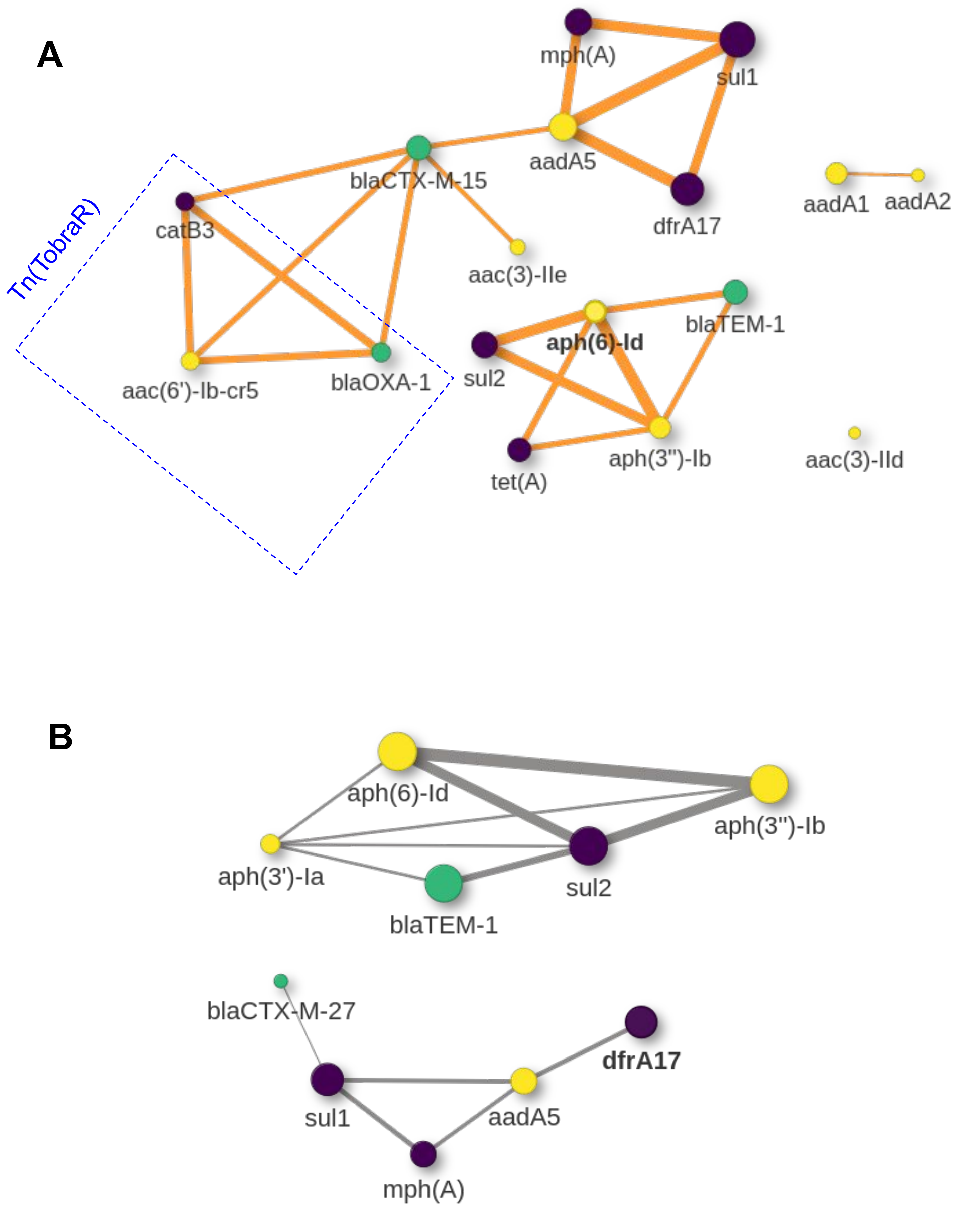


**Supplementary Figure S8. A)** Co-occurrence network of ARGs in the same plasmid prediction in SDD isolates. **B)** Co-occurrence network of ARGs in the same plasmid prediction in baseline isolates. Only connections with a p-value=<0.01 are drawn.
